# Correlation of occludin protein mobility with paracellular leak pathway permeability in renal epithelia

**DOI:** 10.1101/437996

**Authors:** Josephine Axis, Alexander L. Kolb, Robert L. Bacallao, Kurt Amsler

## Abstract

Studies have demonstrated regulation of the epithelial paracellular permeability barrier, the tight junction, by a variety of stimuli. Recent studies have reported a correlation between changes in paracellular permeability, particularly paracellular permeability to large solutes (leak pathway), and mobility of the tight junction protein, occludin, in the plane of the plasma membrane. This had led to the hypothesis that changes in occludin protein mobility are causative for changes in paracellular permeability. Using a renal epithelial cell model system, MDCK, we examined the effect of various manipulations on both leak pathway permeability, monitored as the paracellular movement of a fluorescent molecule (calcein), and occludin protein mobility, monitored through fluorescence recovery after photobleaching. Our results indicate that knockdown of the associated tight junction protein, ZO-1, increases baseline leak pathway permeability, whereas, knockdown of the related tight junction protein, ZO-2, does not alter baseline leak pathway permeability. Knockdown of either ZO-1 or ZO-2 decreases the rate of movement of occludin protein but only knockdown of ZO-2 protein alters the percent of occludin protein that is mobile. Further, treatment with hydrogen peroxide increases leak pathway permeability in wild type MDCK cells and in ZO-2 knockdown MDCK cells but not in ZO-1 knockdown MDCK cells. This treatment decreases the rate of occludin movement in all three cell lines but only alters the mobile fraction of occludin protein in ZO-1 knockdown MDCK cells. Finally, we examined the effect of renal ischemia/reperfusion injury on occludin protein mobility in vivo.

Ischemia/reperfusion injury both increased the rate of occludin mobility and increased the fraction of occludin protein that is mobile. These results indicate that, at least in our cell culture and in vivo model systems, there is no consistent correlation between paracellular leak pathway permeability and occludin protein mobility.

## INTRODUCTION

Until recently, the epithelial cell paracellular permeability barrier, the tight junction, was viewed as a relatively static structure that did not undergo significant regulation. This view has changed dramatically with the demonstration of regulation of the barrier in a variety of epithelia (Rao, 2009; Shen et al., 2011; Steed et al., 2010; Turner et al., 2014). Since then, the mechanism(s) underlying these regulatory events have been under intense investigation. In 2008, Shen et al. (2008) demonstrated that some tight junction proteins exhibit dynamic mobility, moving into and out of the tight junction region. Subsequent studies from several laboratories have demonstrated correlations between a change in the mobility of a tight junction protein and a change in paracellular permeability.

Many studies have focused on the possible role of changes in the mobility of the tight junction membrane protein occludin in mediating regulation of epithelial cell paracellular permeability by various stimuli. Raleigh et al. (2011) reported that phosphorylation of occludin serine 408 by the kinase CK2 in Caco-2 intestinal epithelial cells increased paracellular permeability to ions and small molecules (pore pathway). This phosphorylation also increased the fraction of occludin protein that was mobile within the membrane. Manda et al. (2018) reported a phosphorylation hotspot in the C-terminal region of occludin protein. Deletion of this phosphorylation hotspot both reduced occludin protein mobility and diminished the ability of several stimuli to modulate paracellular permeability. Buschmann et al. (2013) identified a C-terminal region of occludin protein (OCEL domain) that was required for Tumor Necrosis Factor-induced regulation of pore pathway permeability and occludin protein dynamics in Caco-2 intestinal epithelial cells. We previously reported that hydrogen peroxide (H_2_O_2_) both increased renal epithelial cell paracellular permeability to large molecules (leak pathway) and decreased the mobility of full-length occludin protein (Janosevic et al., 2016). The H_2_O_2_-induced regulation of occludin protein dynamics required the C-terminal region of occludin protein. The mutations of occludin protein that diminish regulation of both paracellular permeability and occludin protein dynamics are in regions of the protein believed to be involved in occludin protein binding to ZO proteins, suggesting involvement of this protein-protein binding event in both regulatory responses.

These demonstrated correlations between modulation of occludin protein dynamics and regulation of paracellular permeability have led to the hypothesis that changes in tight junction protein mobility are causative for observed changes in epithelial cell paracellular permeability. We have used the MDCK Type II renal epithelial cell model system and the renal ischemia/reperfusion model in vivo to examine this possible relationship in more detail. Our results argue that changes in occludin protein mobility are neither necessary nor sufficient to produce changes in renal epithelial leak pathway permeability.

## MATERIALS AND METHODS

### Reagents

H_2_O_2_ (3% solution) was obtained from Acros Organics. Calcein was obtained from Invitrogen. α-Modification Minimal Essential Medium was obtained from Corning-CellGro. Heat-inactivated fetal bovine serum was obtained from Atlanta Biologicals. Pencillin/Streptomycin Solution (100X) was obtained from MP Biomedicals. L-Glutamine Solution (200 mM) was obtained from GIBCO Life Technologies. Trypsin/EDTA Solution (0.25%) was obtained from HyClone. Antibodies used in the studies presented here are as follows: rabbit anti-ZO-1 antibody Invitrogen, catalog #40-2200), rabbit anti-ZO-2 antibody (Life Technologies, catalog #711400), HRP-conjugated goat anti-rabbit F_(ab’)2_ fragment antibody (Jackson ImmunoResearch Laboratories, catalog #111-036-003), HRP-conjugated goat anti-rabbit F(_ab’)2_ fragment antibody (Invitrogen, catalog #31461).

### Cell Lines

Wild type MDCK Type II cell line was a gift from Dr. C.M. Van Itallie (NHLBI). The ZO-1 knockdown MDCK Type II cell line and the ZO-2 knockdown MDCK Type II cell line were kind gifts from Dr. Alan Fanning (University of North Carolina). All MDCK Type II cell lines were obtained from the same parental MDCK Type II cell line. Characterization of the ZO-1 knockdown MDCK Type II cell line and ZO-2 knockdown MDCK Type II cell line are described in Medina et al. (2000) and Van Itallie et al. (2009). Cell cultures were checked for mycoplasma contamination upon thaw of a new stock vial and periodically thereafter (MycoAlert Mycoplasma Detection Kit, Lonza). All tests were uniformly negative. New stock cell populations of each cell line were thawed after 15 passages. Cell line authentication was not performed.

Cell populations were grown as stock cultures maintained at a subconfluent density in tissue culture-treated flasks in Complete Medium (αMEM supplemented with 10% fetal bovine serum plus 2 mM L-glutamine plus penicillin/streptomycin) at 37oC in a humidified 5% CO_2_ atmosphere. Cells were passaged every 3-4 days by detaching cells with trypsin/EDTA solution and replating at a 1:10-1:20 dilution onto tissue culture-treated flasks. For flux experiments, detached cells were seeded onto permeable membrane filters (BD Biosciences; 25 mm diameter, 0.4 μm pore diameter) in 6-well tissue culture plates containing 2 ml Complete Medium in both the upper and lower compartments. Medium was replenished every 2-3 days. Twelve-thirteen days after seeding, medium was replenished with serum-free αMEM supplemented with 2 mM L-glutamine plus penicillin/streptomycin. Cell populations were incubated overnight and then used for flux assays as described below and previously (Caswell et al., 2013).

### Paracellular Permeability

For measurement of paracellular permeability, medium was aspirated from both fluid compartments and cell populations were washed 2X with and incubated for 1 hour at 37°C in Ca-Mg-containing phosphate-buffered saline (Ca-Mg-PBS) without or with 55 μM H_2_O_2_ (pretreatment). Basolateral solution was then replaced with Ca-Mg-PBS without or with 55 μM H_2_O_2_ plus 40 μM calcein. Aliquots (20 μl) of the apical compartment fluid were sampled periodically over a 4 hour period. Fluorescence was measured using a Biotek Synergy multiwell plate reader at 495 nm (excitation) and 515 nm (emission) wavelengths. Calcein content in the apical fluid compartment was calculated for each population using a standard curve generated from samples of known calcein content. Data are presented as mean ± standard deviation of triplicate independent samples. Average calcein flux rate for each cell line and condition are expressed as the mean ± standard deviation of at least nine independent experiments.

### Western Blot Analysis

Cell lysates were prepared from cell populations maintained under the conditions used for measurement of paracellular permeability. Protein separation and Western blotting were performed as previously described (Caswell et al., 2013). Briefly, 20 μg protein for each sample was added to separate wells of an 8% polyacrylamide gel and proteins were electrophoretically separated (~120 V). Proteins were electrophoretically transferred to nitrocellulose membrane using an Amersham SemiDry Transfer System at 15 V for 45 minutes. Nitrocellulose membranes were allowed to dry and stored at room temperature. Specific proteins were detected by Western blotting. Nitrocellulose membranes were blocked by incubation with Tris-buffered saline solution plus 0.1% Tween 20 (TBST) containing 3% non-fat dry milk (Blocking Solution) at room temperature for 2 hours with rocking. Nitrocellulose membranes were then incubated in Blocking Solution containing primary antibody for 60 minutes at room temperature with rocking. Membranes were then washed with TBST four times for 5 minutes each and then incubated in Blocking Solution containing peroxidase-conjugated secondary antibody for 60 minutes at room temperature with rocking. Following washing four times for 5 minutes each with TBST, nitrocellulose membranes were incubated in Amersham ECL Prime Western Blotting Reagent (catalog #RPN 2236) according to the manufacturer’s instructions. Chemiluminescence signals were detected using an Amersham RGB800 imager. Primary antibody dilutions for Western blotting were: ZO-1 - 1:500-1:2,000; ZO-2 - 1:500-1:2,000. HRP-conjugated anti-rabbit F_c_ fragment antibodies were used at a dilution of 1:10,000-1:20,000. Presented blots are representative of at least four separate blots obtained from at least four independent sets of samples.

Contents of tight junction proteins in each cell line were normalized to the content of either β- actin or β-tubulin in the same lysate sample. The normalized content of each tight junction protein in the knockdown cell lines was then expressed as a fraction of the normalized content of the same tight junction protein in lysates of wild type MDCK cells. Relative tight junction protein content was measured in at least four independent lysate samples for each cell line. Data are expressed as mean + standard deviation of at least four independent measurements.

### Adenoviral Construct Cloning

Polymerase chain reaction (PCR) was employed to amplify full length hOccludin from the EGFP-hOccludin construct (a gift from Dr. J.R. Turner, Harvard Medical School) and to introduce restriction sites (KpnI and XbaI) using the following primers:

Full-Length hOccludin primers (1,500 bp) -

Forward:

TACAAGTCCGGACTCAGATCTCGAGCTCAAGCTTCGAATTCTGCAGTCGACGGTACCATGTCATCCAGGCCTCTTGA

Reverse:

TGATCAGTTATCTAGATCCGGTGGATCCCGGGCCCGCGGATAAATGTGTTCCTTGTCCCA hOccludin constructs were ligated into the pAcGFP1-Hyg-C1 vector and expanded by transformation into DH5α cells. The full-length hOccludin-EGFP coding sequence was isolated using KpnI and XbaI restriction sites and was amplified by PCR using the following primers: Full-Length hOccludin primers (2,500 bp) Forward: AGATCTCGAGCTCAAGCTTCGAATTCTGCAGTCGACGGTACCATGGTGAGCAAGGGCGAG GAGCTG Reverse: TGATCAGTTATCTAGATCCGGTGGATCCCGGGCCCGCGGATAAATGTGTTCCTTGTCCCA Full-length hOccludin-EGFP coding sequence was obtained using the KpnI and XbaI restriction sites and was cloned into the pAdEasy-1 adenoviral vector according to the manufacturer’s instructions (Agilent Technologies, catalog# 240010-12).

### Fluorescence Recovery After Photobleaching Cell Culture

Cell populations seeded onto sterile 18 mm round cover glass pieces placed in sterile 35 mm petri dishes. Cell populations were grown to and maintained at confluence for at least 7 days. Cell populations were transduced with an adenovirus containing full-length EGFP-hOccludin construct (pAdenovirus-hOccludin-FL-EGFP) at a 50:1 MOI. Transduced cells were serum starved overnight (αMEM, 2 mM L-glutamine, penicillin/streptomycin). FRAP experiments were performed 3-6 days after viral transduction. Transduced cell populations were treated with Ca-Mg-containing PBS ± 55 μM H_2_O_2_ for 2 hours at 37°C. Cells were then analyzed by Fluorescence Recovery After Photobleaching (FRAP). Cover slips containing cell populations were mounted onto a chamber and incubated in Ca-Mg-PBS maintained at 37°C. Cells were imaged using a Leica TCS SP5 inverted microscope with 63.0 × NA 1.40 oil-immersion objective. A Region of Interest (ROI) at approximately the center of the area to be photobleached (comprising less than 50% of the total photobleached area) was designated. Data were collected and processed using Leica Microsystems LAS AF software. The settings for the FRAP procedure were: Argon laser power = 5%; Leica-488 nm laser power = 5%-10%; pre-bleach - 5 frames per 1.318 second; bleach - 5 frames; post-bleach - 1-5 frames per 1.318 seconds; post-bleach - 2-20 frames per 5 seconds; post-bleach - 3-40 frames per 10 seconds.

#### In Vivo: Confocal Microscopy Setup and Intravital Imaging

Rats were anesthetized with an intraperitoneal injection of 130 mg/kg of thiobutabarbital (Inactin Hydrate). Animal respiration, reflex response, color, and body temperature were monitored throughout the surgical procedure. Once the animal was fully anesthetized, the left flank was shaved and then the animal’s left kidney was located. Following location, a small incision was made on the flank and the kidney was exposed through the incision. The animal was then transported to the imaging room for renal imaging. The kidney was placed face down into a 35 mm glass dish allowing the rat bodyweight to stabilize the interaction between the kidney and the stage to reduce motion. Saline was poured into the dish to ensure the kidney did not dry out. An Olympus FV 1000 MPE microscope (Olympus, Tokyo, Japan) with a Spectra Physics MaiTai Deep Sea Laser (710-990 nm) with a 20X or 60X water immersion objective was used for imaging. Intravital images were acquired using wavelengths ranging from 820-860 nm. This system is also equipped with two external detectors for multi-photon imaging and dichroic mirrors to collect red, green, and blue light emissions. This light is collected via a three-band pass filter system. Blue light is collected between 420-460 nm, green light between 495-540 nm, and red light between 575-630 nm. For EGFP studies, the pseudo-green and pseudo-red channels were merged to eliminate the possibility of tubule autofluorescence.

### In Vivo: FRAP

Four days prior to imaging male Sprague-Dawley rats (250-350g) received hydrodynamic delivery of adenoviral EGFP-occludin or EGFP-occludin 860 as previously described (Corridon et al., 2013). Animals were prepared for intravital light microscopy as previously described. For FRAP, EGFP-occludin images were acquired in multiphoton mode and bleaching of the EGFP-occludin protein occurred in one photon mode. Our FRAP studies followed the methods established by Zheng et al. (Zheng et al., 2011).

This study was carried out in strict accordance with the recommendations in the Guide for the Care and Use of Laboratory Animals of the National Institutes of Health. The protocol was approved by the Indiana University Institutional Animal Care and Use Committee (IACUC). Injections were performed under Isoflurane and buprenorphine analgesia and all imaging was performed under Inactin analgesia. All efforts were made to minimize suffering.

## RESULTS

### Effect of Knockdown of ZO-1 Protein Versus ZO-2 Protein

We first examined the effect of knockdown of ZO-1 protein versus ZO-2 protein on calcein flux rate and full-length EGFP-occludin protein mobility. Knockdown of ZO-1 protein and ZO-2 protein was confirmed by Western blot analysis (Figure 1a). Compared to wild type MDCK cells, ZO-1 knockdown (ZO-1 KD) cells express 7 ± 5% ZO-1 protein (n=4) and 119 ± 54% ZO-2 protein (n=4). Compared to wild type MDCK cells, ZO-2 knockdown (ZO-2 KD) cells express 98 ± 60% ZO-1 protein (n=4) and 0 ± 0% ZO-2 protein (n=4). Leak pathway permeability was monitored as the paracellular flux of the fluorescent molecule calcein (Caswell et al., 2012). We have previously shown that calcein flux rate is a valid measure of paracellular leak pathway permeability in MDCK Type II cells. As we previously reported, MDCK Type II cells in which ZO-1 protein was knocked down exhibited increased calcein flux rate compared to wild type MDCK Type II cells (p<0.0001; Figure 1b; Table 1; Bilal et al., 2018). MDCK Type II cells in which ZO-2 protein was knocked down, in contrast, exhibited a similar calcein flux rate to wild type MDCK cells (p=0.905; Figure 1b; Table 1).

**Table 1.** Effect of H_2_O_2_ on calcein flux and FRAP parameters in wild type MDCK Type II cells and in MDCK Type II cells in which either ZO-1 or ZO-2 protein has been knocked down.

| Cell Line | Treatment | Calcein Flux<br>(pmoles calcein/cm<sup>2</sup>) | $t_{1/2}$<br>(seconds) | Immobile Fraction (%) |
| --- | --- | --- | --- | --- |
| MDCK | 0 μM $H_2O_2$ | $5.61 \pm 1.62$<br>(n=30) | $50.18 \pm 10.83$<br>(n=21) | $45 \pm 21$ (n=21) |
| | 55 μM $H_2O_2$ | $14.06 \pm 7.48$<br>(n=29) | $80.98 \pm 14.14$<br>(n=18) | $40 \pm 17$ (n=18) |
| ZO-1 KD | 0 μM $H_2O_2$ | $15.11 \pm 4.82$<br>(n=21) | $43.18 \pm 7.23$<br>(n=14) | $44 \pm 14$ (n=14) |
| | 55 μM $H_2O_2$ | $14.44 \pm 5.09$<br>(n=21) | $68.72 \pm 14.85$<br>(n=25) | $29 \pm 13$ (n=25) |
| ZO-2 KD | 0 μM $H_2O_2$ | $5.69 \pm 1.71$<br>(n=9) | $73.86 \pm 15.00$<br>(n=27) | $26 \pm 11$ (n=27) |
| | 55 μM $H_2O_2$ | $12.15 \pm 5.45$<br>(n=9) | $85.22 \pm 14.48$<br>(n=23) | $24 \pm 16$ (n=23) |

Calcein flux data are expressed as the mean ± standard deviation of the indicated number of independent calcein flux experiments. Fluorescence recovery halftime and immobile fraction data, calculated using the Leica LAS AF software, are presented as the mean ± standard deviation of the indicated number of independent FRAP experiments.

**Figure 1.**
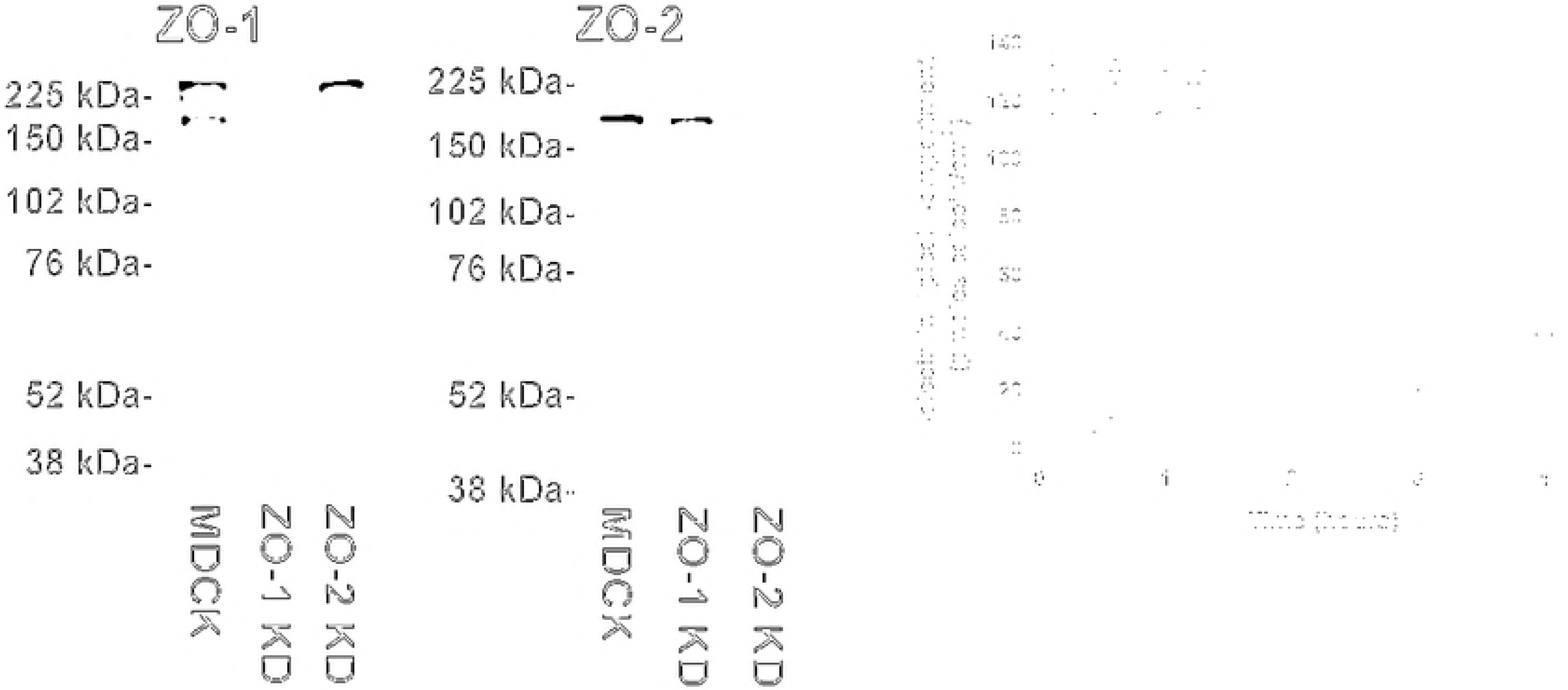
ZO protein content and calcein flux rate in wild type and knockdown MDCK Type II cell lines. (a) Cellular content of ZO-1 protein (left-side blot) and ZO-2 protein (right-side blot) in wild type MDCK cells (MDCK), ZO-1 knockdown MDCK cells (ZO-1 KD), and ZO-2 knockdown MDCK cells (ZO-2 KD). The presented figure is representative of four separate experiments performed using four independently-derived cell lysates.(b) Time course of calcein paracellular flux across post-confluent monolayers of wild type MDCK cells (wild type MDCK), ZO-1 knockdown MDCK cells (ZO-1 KD MDCK), and ZO-2 knockdown MDCK cells (ZO-2 KD MDCK). Each data point represents the mean + standard deviation of three independent samples.

We next determined the mobility of full-length EGFP-occludin protein in these three cell lines. To determine occludin protein dynamics, MDCK Type II cells were transduced with viral expression vectors encoding full-length Enhanced Green Fluorescence Protein (EGFP)-occludin protein. Fluorescence in a region of the tight junction was photobleached and recovery of fluorescence into the photobleached region was monitored over time. Occludin protein dynamics were defined by two parameters: 1) fluorescence recovery halftime (t_1/2_) - the time required to reach half of maximal fluorescence intensity following photobleaching; and 2) immobile fraction - the percentage of the expressed EGFP-occludin which cannot diffuse into the photobleached area during recovery.

In wild type MDCK Type II cells, a portion of full-length EGFP-occludin protein migrated into the photobleached tight junction region over time. The halftime for fluorescence recovery was 50.18 ± 10.83 seconds (Table 1). The immobile fraction of expressed full-length EGFP-occludin protein was 45 ± 21%. Full-length EGFP-occludin expressed in ZO-1 knockdown MDCK Type II cells exhibited a statistically significant increase in fluorescence recovery halftime (p=0.028) but no change in immobile fraction (p=0.946) compared to the parameters measured in wild type MDCK Type II cells. Full-length EGFP-occludin protein expressed in ZO-2 knockdown MDCK cells exhibited a statistically significant increase in halftime (p<0.0001) and a statistically significant decrease in immobile fraction (p=0.0008) compared to the parameters measured in wild type MDCK Type II cells (Table 1).

### Effect of H_2_O_2_ Treatment

We next examined the effect of treatment with sublethal concentrations of H_2_O_2_ on leak pathway permeability and EGFP-occludin protein mobility in these three cell lines. As reported previously (Janosevic et al., 2016), treatment of wild type MDCK cells with sublethal concentrations of H_2_O_2_ increased calcein flux rate (Figure 2a; p<0.0001). Similar to wild type MDCK cells, treatment of ZO-2 knockdown MDCK cells with H_2_O_2_ increased calcein flux rate compared to wild type MDCK cells (Figure 2c; p=0.004). In contrast, treatment of ZO-1 knockdown MDCK cells with H_2_O_2_ did not increase calcein flux rate compared to wild type MDCK cells (Figure 2b; p=0.660).

**Figure 2.**
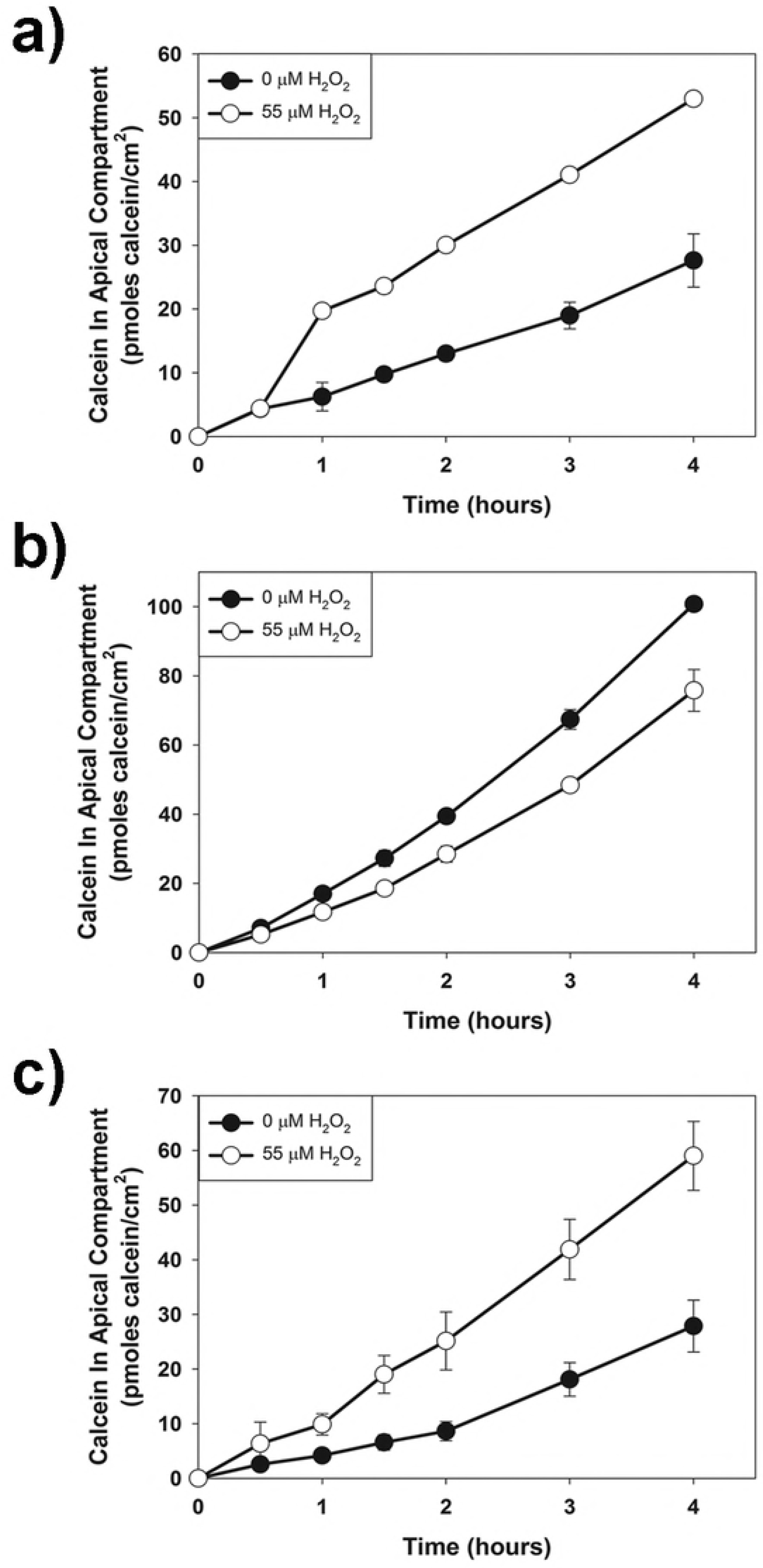
Effect of H_2_O_2_ treatment on calcein flux rate across wild type MDCK Type II cells (a), ZO-1 knockdown MDCK Type II cells (b), and ZO-2 knockdown MDCK Type II cells (c) Post-confluent cell populations were pretreated with 55 μM H_2_O_2_ for 1 hour prior to initiation of the calcein flux assay. Each data point represents the mean ± standard deviation of triplicate independent samples.

As reported previously, treatment of wild type MDCK cells with H_2_O_2_ increased fluorescence recovery halftime (p<0.0001) but did not alter the immobile fraction (p=0.437) compared to untreated wild type MDCK cells (Table 1). Treatment of ZO-1 knockdown MDCK cells with H_2_O_2_ increased fluorescence recovery halftime (p<0.0001) and also decreased the immobile fraction (p=0.003) compared to untreated ZO-1 knockdown MDCK cells. Treatment of ZO-2 knockdown MDCK cells with H_2_O_2_ increased fluorescence recovery halftime (p=0.009) but did not affect the immobile fraction (p=0.623).

### Effect of Renal Ischemia/Reperfusion Injury In Vivo

Finally, we investigated the effect of renal ischemia/reperfusion injury on occludin protein mobility *in vivo* in rats. One of the consequences of renal ischemia/reperfusion injury is an increase in the backflux of forming urine due to a breakdown of the paracellular permeability barrier (see e.g., Kwon et al., 1998). Full-length EGFP-occludin construct in a viral expression construct was introduced into rat renal epithelial cells by retrograde pressure transduction (Corridon et al., 2013). Renal ischemia/reperfusion injury was produced by the renal single clamp model (Kolb et al., 2018). The unclamped kidney was used as a control. In the absence of renal clamp, virtually all of the expressed EGFP-occludin was immobile (92 ± 10%, n=4) and the fluorescence recovery halftime was greater than 500 seconds (n=4). Following ischemia/reperfusion injury, the immobile fraction decreased to 32 ± 15% (n=4; p<0.003 compared to pre-ischemia condition) and the fluorescence recovery halftime decreased to 101 ± 26 seconds (n=4). Because we were unable to measure a fluorescence recovery halftime in the pre-ischemia condition, we were unable to calculate a p value comparing the pre-and post-ischemia conditions for this parameter. However, the data point to a significant difference in baseline dynamics of tight junction protein mobility in vivo as compare to in vitro cell culture studies.

## DISCUSSION

A summary of our results is shown in Table 2. Our results do not demonstrate a consistent correlation between any change in occludin protein dynamics and an increase in paracellular leak pathway permeability. Further, our previous results suggest no consistent correlation between changes in pore pathway permeability and changes in occludin protein dynamics (Janosevic et al., 2016). Our results indicating the lack of a strong correlation between occludin protein dynamics and changes in paracellular permeability in renal epithelia is consistent with the findings of Van Itallie et al. (2016). They reported that knockout of the protein TOCA-1 in MDCK Type II renal epithelial cells increased leak pathway permeability without altering pore pathway permeability. They did not detect any change in occludin protein dynamics associated with this change in leak pathway permeability.

**Table 2.** Summary of changes in paracellular flux rate (Leak pathway) and EGFP-occludin FRAP parameters.

| Cell Type | Manipulation | Flux | $t_{1/2}$ | Immobile Fraction |
| --- | --- | --- | --- | --- |
| MDCK | + H<sub>2</sub>O<sub>2</sub> | ↑ | ↑ | NC |
| ZO-1 KD | + H<sub>2</sub>O<sub>2</sub> | NC | ↑ | ↓ |
| ZO-2 KD | + H<sub>2</sub>O<sub>2</sub> | ↑ | ↑ | NC |
| ZO-1 KD | Compared to MDCK | ↑ | ↑ | NC |
| ZO-2 KD | Compared to MDCK | NC | ↑ | ↓ |
| Renal Epithelia In Vivo | Ischemia/Reperfusion Injury | ↑ | ↓ | ↓ |

↑ - Increase
↓ - Decrease
NC – No change

As mentioned above, binding of occludin protein to ZO proteins is believed to be an important regulator of occludin protein mobility. We have shown that deletion of the ZO binding site in occludin protein increases the fraction of occludin protein that is mobile (data not shown). A second finding of our study is that the presence of ZO-1 versus ZO-2 protein differentially affects occludin protein mobility in renal epithelial cells in culture. In both ZO-1 knockdown MDCK Type II cells and ZO-2 knockdown MDCK Type II cells, the halftime for fluorescence recovery of EGFP-occludin protein was slowed compared to its recovery in wild type MDCK Type II cells. This suggests that the presence of ZO-1 versus ZO-2 protein does not differentially affect the rate of movement of mobile EGFP-occludin molecules. In contrast, the fraction of the EGFP-occludin protein that was immobile was decreased in ZO-2 knockdown cells but was unaffected in ZO-1 knockdown cells compared to wild type MDCK cells. This suggests that occludin protein anchoring, at least for EGFP-occludin moving into the tight junction region, is mediated preferentially through binding to ZO-2 protein.

A third finding of our study is that there is no clear correspondence between the dynamic behavior of full-length occludin protein in renal epithelial cells in culture versus in renal epithelial cells *in vivo*. Occludin protein expressed in renal epithelial cells *in vivo* was virtually immobile, whereas, occludin protein expressed in renal epithelial cells in culture exhibited a substantial mobile fraction. Further, there was little correspondence between the H_2_O_2_-induced changes in occludin protein dynamics in renal epithelial cells in culture and the effects of ischemia/reperfusion injury on occludin protein dynamics in kidney epithelial cells *in vivo*. A caveat to this conclusion is that H_2_O_2_ treatment of cells in culture may not be a complete model for the effects of renal ischemia/reperfusion injury *in vivo*. But it does point to the need for caution when extrapolating findings with cultured cells to the *in vivo* environment.

In summary, our results indicate that changes in occludin protein dynamics are neither necessary nor sufficient to induce changes in leak pathway permeability in the MDCK Type II renal epithelial cell system. It is possible that our results are relevant only to our MDCK Type II renal epithelial cell model system. We cannot rule out the possibility that other epithelial cell types exhibit different behaviors. It is also possible that changes in pore pathway permeability may be more strongly correlated with changes in occludin protein dynamics. Since our model system did not exhibit changes in pore pathway permeability using our manipulations, we were unable to assess this possible relationship. Previous studies, however, reported that the ZO-2 knockdown MDCK Type II cells used in our study did not exhibit any change in pore pathway permeability compared to wild type MDCK Type II cells (Van Itallie et al., 2009). Despite this lack of an effect on pore pathway permeability, our data demonstrate substantial changes in occludin protein dynamics in ZO-2 knockdown MDCK cells compared to wild type MDCK cells arguing against this correlation.

